# RGG-motif protein Scd6 affects oxidative stress response by regulating Cytosolic caTalase T1 (Ctt1)

**DOI:** 10.1101/2023.09.05.556329

**Authors:** Sweta Tiwari, Sudharshan Sj, Purusharth I Rajyaguru

**Affiliations:** Indian Institute of Science, Bengaluru; Duke University; Indian Institute of Science

**Keywords:** Translation, Post-transcriptional control, H_2_O_2,_ Catalase, Scd6, Reactive oxygen species (ROS), oxidative stress, RGG-motif proteins, stress response

## Abstract

In response to stress, cells undergo gene expression reprogramming to cope with external stimuli. As translation is energy consuming process, its regulation during stress is crucial for cellular adaptation. Cells utilize a conserved stress response mechanism called global downregulation of translation, leading to the storage of translationally repressed mRNAs in RNA granules or RNP condensates. During oxidative stress induced by H_2_O_2_, genes responsible for combating oxidative stress, such as catalases and glutathione peroxidase, are strongly induced. However, the post-transcriptional regulatory events affecting these genes during H_2_O_2_ stress are not well-explored. RNA binding proteins such as RGG motif proteins play a critical role in mediating translation regulation and have diverse physiological functions. Scd6, an RGG motif-containing protein in yeast, acts as a translational repressor through its interaction with eIF4G. Although the role of Scd6 as a translational repressor and decapping activator is known, its specific mRNA targets are largely unknown.This study identifies the role of Scd6 in oxidative stress response by regulating cytoplasmic catalaseT1 (*CTT1)*. Altering Scd6 levels influenced Ctt1 protein levels, thereby affecting oxidative stress response. Scd6 overexpression increased sensitivity, while deletion enhanced tolerance to H_2_O_2_ treatment due to ROS accumulation in the yeast cell. In response to H_2_O_2_ treatment, Scd6 forms dynamic puncta which contains RNA. Overall, this study proposes regulation of oxidative stress response via modulation of *CTT1* mRNA by Scd6.

## Introduction

In stress conditions, the reprogramming of the gene expression is a must for a cell to respond to the external stimuli. As translation is a highly complex and energy expensive process, its regulation under stress conditions becomes crucial for the cell. Global downregulation of translation is a conserved stress response mechanism employed by cells to re-orient their gene expression programs^1,2^.Translationally repressed mRNAs are often stored in RNA-protein complexes known as RNA granules or RNP condensates^3^ which are conserved and dynamic structures. Upon various stresses like glucose starvation, heat shock, nitrogen starvation, cells form membraneless cytoplasmic bodies like stress granules or P bodies^4–6^. Cytoplasmic RNA granules could be sites of mRNA storage and degradation^7^.

Upon H_2_O_2_ induced oxidative stress, there is strong induction of genes involved in combating oxidative stress such as catalases^8^ and glutathione peroxidase. These enzymes can neutralize H_2_O_2_^9^. While previous studies have established correlations between transcriptional changes and H_2_O_2_ stress^2,10–12^, the post-transcriptional regulatory events remain poorly explored.Catalase is a heme containing antioxidant enzyme induced in response to oxidative stress,heat shock and starvation^13,14^. In yeast, the cytoplasmic catalase is coded by *CTT1* gene and the peroxisomal catalase is coded by *CTA1* gene^15^. Transcriptional regulation of *CTT1* is well-explored.It is regulated by proteins such as Msn2p/Msn4p, Hog1p, Hap1p, Yap1p, and Zap1p acting upon upstream activating elements (UAS) in the *CTT1* promoter^14,16,17^.

RNA binding proteins play a key role in mediating various steps required for the regulation of translation. An emerging and second largest class of RNA binding proteins is RGG motif proteins^18,19^. These proteins are characterized by the presence of Glycine and Arginine repeats, also termed RGG /RG Box^19^. RGG motif proteins play functional roles in various physiological processes such as transcription, pre-mRNA splicing, DNA damage signaling, mRNA translation, and the regulation of apoptosis. RGG motif apart from interacting with RNA can also interact with proteins^19^. Often the RGG-motifs are fused to canonical RNA and protein interaction domains which enhance their ability to regulate mRNA fate ^20^.

Scd6, an RGG motif-containing protein in *S. cerevisiae*, represses translation by preventing the formation of the 48S pre-initiation complex by binding to eIF4G via its RGG motif^21^. However, the integrity of the cap-binding complex is not lost during repression which suggests that the repressed mRNA can re-enter translation upon specific physiological cues. Arginine methylation of the RGG-motif augments the interaction of Scd6 with eIF4G which enhances the translation repression^22^. Scd6 binds itself in RGG-motif dependent manner and self-association regulates its repression activity^23^. Although Scd6 has important roles as a translational repressor and an indirect decapping activator, detailed analysis of specific mRNA targets remains unknown.Interestingly, RNA-Seq analysis of polysome-associated mRNAs in *Δscd6* strain suggests increased translational efficiency of *CTT1 mRNA* in Δ*scd6* background^24^.

In this study, we explored the role of Scd6 in oxidative stress response by translational regulation of *CTT1*.We observe that, overexpression and deletion of Scd6 led to changes in Ctt1 protein levels without affecting mRNA levels. Notably, altered Ctt1 protein levels influence the cellular response to oxidative stress, with Scd6 overexpression making cells more sensitive and Scd6 deletion rendering cells more tolerant to H_2_O_2_ treatment. This sensitivity and tolerance were found to be a consequence of ROS accumulation upon H_2_O_2_ stress. In response to H_2_O_2_ treatment, Scd6 forms dynamic puncta which contains RNA. We further observe that, in the absence of any stress, *CTT1* remains in association with Scd6 as an mRNA target. Putting together, we propose a mechanism for the translational regulation of *CTT1* by Scd6.

## Materials and methods

### Yeast strains and plasmids

Yeast strains used in this study is BY4741 and its derivatives (Listed in table 1). These strains were grown either in yeast extract/peptone media or synthetic complete minimal media at 30 °C at 220 rpm.

**Table no.1:** The strains used in the study.

| Strain | Description |
| --- | --- |
| Wild type | MATa his3Δ1 leu2 met15 ura3 (BY4741) |
| Δscd6 | MATa his3Δ1 leu2 ura3 his3 met15 *scd6*Δ::KanMX |
| Ctt1-GFP | MATa his3Δ1 leu2 met15 ura3 Ctt1-GFP |
| Ctt1-GFP Δscd6 | MATa his3Δ1 leu2 met15 ura3 Ctt1-GFP *scd6*Δ::KanMX |
| Pab1RFP | A gift from Prof. Beidong Liu's Lab |

**Table no.2:** List of the plasmids used in the study.

|  |  |
| --- | --- |
| pYES EV | 2μ empty vector, Ura and Amp<sup>R</sup> selection marker |
| pYES Scd6GST | Expressing Scd6GST under its own promoter |
| pRS316 Scd6GFP | CEN plasmid expressing Scd6GFP, Ura and Amp<sup>R</sup> selection marker |
| pGP EV | pGP564, 2μ empty vector, Leu- selection marker |
| pGP Scd6GFP | pGP564, Expressing Scd6GFP under its own promoter |
| pRS416 Edc3mch | Expressing Edc3mch, Ura selection marker |

**Table no.3:** List of the antibodies used in the study.

| Antibody | Description |
| --- | --- |
| α-GFP | BioLegend, Catalog no. 338001 |
| α-mono methyl arginine | CST, catalog no. 8711 |
| α-PGK1 | abcam, Catalog no. ab113687 |
| α-Mouse | Jackson ImmunoResearch Laboratories, Code No. 115- 035-003 |
| α-Rabbit | Jackson ImmunoResearch Laboratories, Code No. 111- 035-003 |
| α-GST | CST 26H1 Mouse mAb Catalog no. 2624 |

### Yeast transformation

WT cells were diluted from an overnight grown primary culture to 0.1 OD_600_ and grown till 0.6 OD_600._ These cells were then pelleted down at 5000 rpm. The pellets were washed with water followed by 100mM LiAc (2 times). After the washing step, the cells were resuspended using 300μL of 100mM LiAc and further aliquoted into 3 different microcentrifuge tubes. To the cell suspension, 240μL of 50 % PEG (v/v), 36μLof 1 M LiAc, and 25μL of salmon sperm DNA (100 mg/ml) were added one by one and vortexed vigorously for a few seconds. To the control, no template DNA was added. The mixture was then incubated at 30°C for 30 minutes, followed by 15 minutes of heat shock at 42°C. The cells were then pelleted. The pellet was resuspended using 100 μL of H_2_O and plates on SD Ura-/SD Leu-plates and followed by incubation at 30°C for 2 days^25^.

### H_2_O_2_ treatment

Yeast cells were diluted to an OD_600_ of 0.1 from an overnight grown primary culture and allowed to grow till 0.4-0.8 OD. These cells were then treated 4mM H_2_O_2_ for 30 minutes at 30 °C at 220 rpm in dark^8,26^. Cells were then pelleted at 4200rpm for 1 minute at RT and stored in -80°C till further use.

### Colony-forming unit (CFU) assay

The yeast cells were diluted at an OD_600_ of 0.1 from an overnight grown primary culture and allowed to reach 0.8 OD_600_. After it reached 0.8 OD_600_, the cultures were split into two parts: to one part, 4mM H_2_O_2_ was added, and an equal amount of water was added to the other half of the culture. Both were allowed to incubate at 30°C for 30 minutes at 220 rpm. The cells were then serially diluted, followed by plating two different dilutions (0.001 and 0.0005) on Synthetic Complete Minimal media agar plates. The plates were incubated at 30 °C for 30 minutes at 220 rpm. The CFU count was used to determine the survival^26^. The percentage survival was normalized to the untreated condition. Statistical analysis was done using GraphPad Prism Version 8.0. Statistical significance was calculated using a two-tailed paired t-test.

### Detection of Intracellular Reactive Oxygen Species (ROS) using H_2_DCFDA staining

Intracellular ROS levels were monitored using 2,7-dichlorofluorescein diacetate (H_2_DCFDA) staining. WT cells with Δscd6/ Scd6GST of 0.8OD_600_ were treated with 4mM H_2_O_2_ for 30 minutes at 30 °C followed by washing with 1x Phosphate Buffer Saline (PBS). These cells were incubated with 10μM H_2_DCFDA^27^ for 15 minutes followed by washing with 1x PBS. After resuspending the cells with 1x PBS, fluorescence was measured at the excitation wavelength of 504nm and emission wavelength of 524 nm using Tecan infinite pro 500.

### Cycloheximide treatment

Yeast cells were diluted to an OD_600_ of 0.1 in 10ml SD Ura-with 2% glucose from an overnight grown primary culture and allowed to grow till 0.4 -0.5 OD_600_. These cultures were divided into two parts, 5ml culture was treated with 4mM H_2_O_2_ (2.04μl of 9.8M stock) and same amount of H_2_O was added into another 5ml culture (untreated) followed by 30 minutes incubation at 30 °C at 220rpm under dark. The H_2_O_2_ treated culture was washed off using SD Ura-. To the 5ml of H_2_O_2_ treated culture, either 5μl of 100μg/ml cyloheximide^6,28^ or 5μl methanol (control) was added. Cells were incubated for 10 minutes at 30 °C at 220 rpm under dark. These cells were then pelleted and taken for live cell imaging.

### Yeast live cell imaging

Post-H_2_O_2_/cycloheximide treatment, the cells were pelleted at 14000 rpm for 15 seconds, spotted on glass cover slip and observed using live cell imaging. Yeast images were acquired using a Deltavision RT microscope system running softWoRx 3.5.1 software (Applied Precision, LLC), using an Olympus 100×, oil immersion 1.4 NA objective. The Green Fluorescent Protein (GFP) channel had 0.5 seconds of exposure and 50 % transmittance. The Red fluorescent channel (mcherry) had 0.6 seconds of exposure and 50% transmittance. Granules per cell were counted for minimum 100 cells for each experiment. Statistical analysis was done using GraphPad Prism Version 8.0. Statistical significance was calculated using two tailed unpaired/paired t-test.

### Pulldowns and western blotting

For gluthathione pull-downs^22^, 150ml culture pellets of yeast cells were split into two tubes and lysed using beat beating for 30 minutes. The lysis buffer contained 10mM Tris (pH 7.5), 150mM NaCl, 0.5mM EDTA and 0.1% NP40. Post-bead beating, cells were spun at 5500rpm for 10 minutes at 4°C. The dilution buffer (10mM Tris (pH 7.5), 150mM NaCl, and 0.5 mM EDTA) was added to the supernatant. The input was collected from the mix and 50μl of equilibrated glutathione sepharose 4B (GE healthcare) were added to the reaction mix. The reaction mix was nutated for 2 hours at 4°C followed by spin at 1500rpm for 30 seconds. The beads were washed 2 times using the dilution buffer. The beads were finally resuspended with 200μl of dilution buffer. 2.5% of Input and 5% PD was loaded on 8% SDS-PAGE gel. Rest of the pulldown fraction was used for RNA isolation.

For checking the methylation status of Scd6, pGP564 EV and pGP564 Scd6GFP were transformed in BY4741. 7.5μl of equilibrated magnetic GFP trap beads (ChromoTek) were added to the reaction mix. Rest of the procedure in GFP trap pulldown was same as described in the glutathione pulldown.mono methyl arginine (MMA) antibody (Cell Signaling Technology) CST, catalog no. 8711; 1:1000 dilution) was used for detecting the methylation signal of Scd6 with α-rabbit as secondary antibody.

### RNA isolation

RNA isolation was performed using the hot acidic phenol method^29^. The mid-log phase cells were harvested by centrifugation. The cell pellet was resuspended using DEPC treated autoclaved MQ. The cells were again collected by a flash spin at 10k rpm. The cell pellet was re-suspended with 400 μL of TES solution (10mM Tris Chloride pH 7.5, 10mM EDTA and 0.5% SDS). 400 μLof hot acidic phenol was added to the tube, followed by vigorous vortexing for 10 seconds. The tubes were incubated at 65°C for 60 minutes with vortexing at every 15 minutes. After incubation, it was kept on ice for 5 minutes followed by spinning at 14000rpm 10 minutes, 4 °C. The aqueous layer obtained by this step was transferred into another tube already having 400 μL of Chloroform. It was vortexed vigorously for 10 seconds. The tubes were spun at 14000 rpm for 10 minutes, 4 °C. The aqueous layer was transferred into another tube carefully. To this tube, 1/10th vol of 3M NaAc pH 5.2 and 2.5 vol of EtOH was added. This mixture was snap-chilled using liquid N2. The tubes were spun at 14000 rpm for 10 minutes, 4 °C. The pellet was washed twice using 70% EtOH, air-dried, and resuspended using 100 μL DEPC MQ. RNA quality was checked by 1% agarose formamide gel electrophoresis.

### Reverse transcription and quantitative real-time PCR

The isolated RNA was treated with DNaseI (Thermo, EN0525). 5 μg of total RNA, 2.5 units of DNaseI, and DNaseI Buffer with MgCl_2_ were added for a 30 μl reaction. The above mixture was incubated at 37°C for 30 minutes. After incubation, 3 μl of 50mM EDTA was added to stop the DNAaseI reaction. The DNAaseI-treated RNA was checked using 1.2 % agarose formamide gel electrophoresis to assess RNA quality. DNaseI treated RNA was then used for cDNA synthesis. According to the manufacturer’s protocol, one microgram of RNA was used to synthesize cDNA using the RevertAid RT Reverse Transcription Kit (Thermo, K1691). cDNA was diluted at 1:10, and real-time PCR was performed using TB Green™ Premix Ex Taq™ (TaKaRa). For qRT-PCR, three technical replicates were assembled with 2ul cDNA/reaction and 0.5μM each primer in a BioRad iQ5 Real-Time PCR Detection System. The PCR conditions were 95° C for 12 min for initial denaturation, followed by 30 cycles of 95°C for 20 sec, 46° C for 30 sec, and 72° C for 30 seconds. DNA was quantified in every cycle at the extension step. Melt curve acquisition was carried out at 64°C for 8 sec. Ct values were extracted with auto baseline and manual threshold. ΔΔCt method was used to calculate the final log2 FoldChange values, which were then plotted on a 19 box and whisker plot using GraphPad prism 7.0. Significance was calculated by a ratio unpaired t-test.

## Results

### Scd6 affects Ctt1 protein levels

We first tested the effect of Scd6 overexpression on Ctt1 protein levels in the untreated and 4mM H_2_O_2_ treated mid-log phase cells. In mid-log phase untreated cells, we observed that Ctt1 protein levels increased upon Scd6 overexpression (lane 1, 2) (Figure 1a). We further observed a decrease in Ctt1 protein levels upon Scd6 overexpression in mid-log phase cells treated with 4mM H_2_O_2_ (lane 3, 4) (Figure 1a). There was an induction of Ctt1 protein levels in wild-type cells (transformed with empty vector – EV) upon H_2_O_2_ treatment during the mid-log phase (Figure 1a).The H_2_O_2_ treatment in our experiment is working since we observe induction of Ctt1 protein upon H_2_O_2_ exposure(lane 1 and 3) as reported earlier^8^. There was no significant difference observed in the *CTT1* mRNA levels in EV and Scd6GST (FigureS1 a, b).

**Figure 1:**
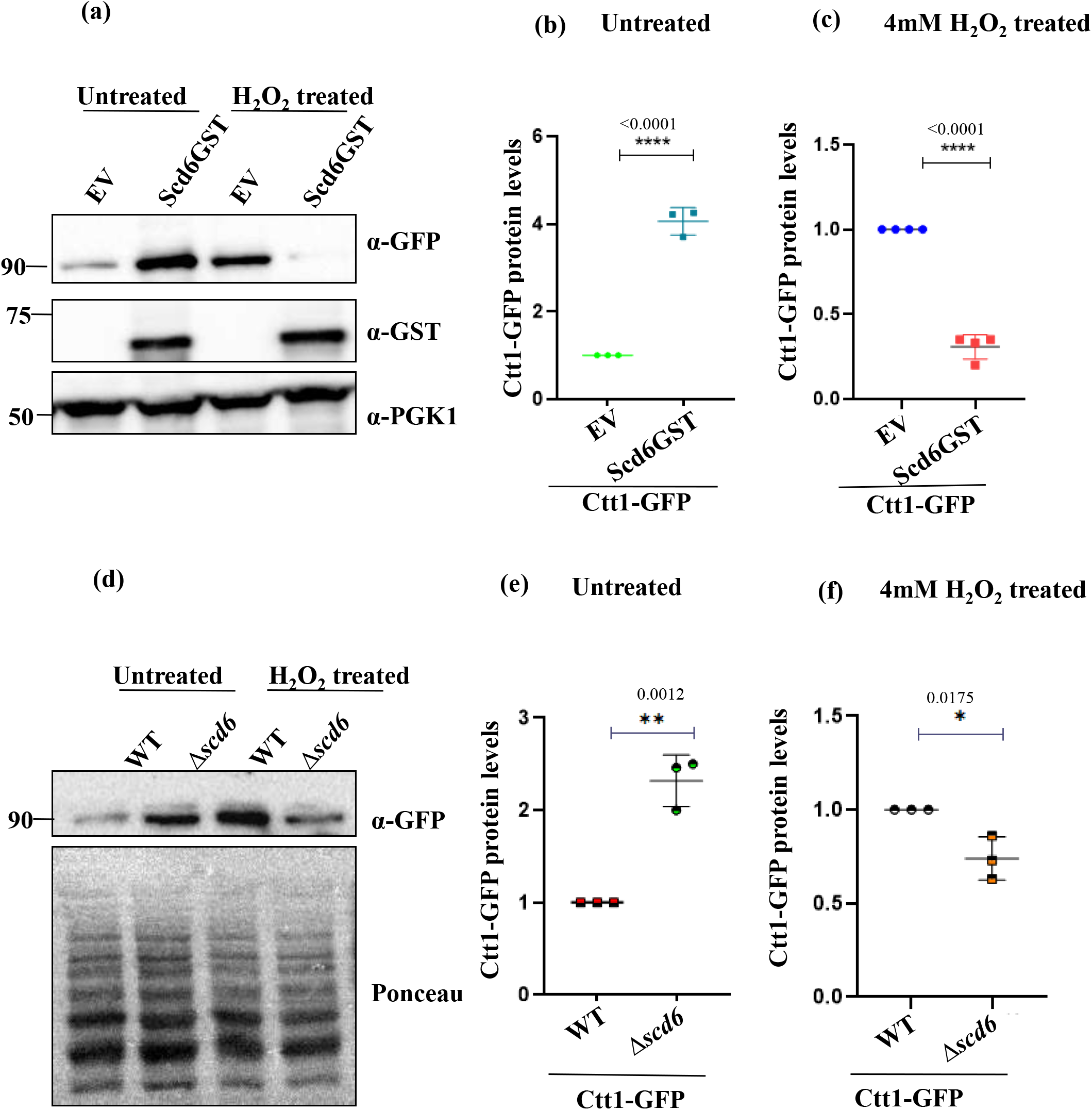
Scd6 affects Ctt1 levels: (a) Ctt1-GFP protein levels upon Scd6 overexpression under untreated and treated condition (4mM H_2_O_2_) Quantification for (b) untreated (c) 4mM H_2_O_2_ treated condition. PGK1 was used as a loading control (d) Ctt1-GFP protein levels in WT and *Δscd6* under untreated and treated condition (4mM H_2_O_2_) Quantification of Ctt1-GFP protein levels under (e) untreated and (f) 4mM H_2_O_2_ treated condition. The GFP signal was detected by α-GFP antibody. Ponceau was used as a loading control. Statistical significance was calculated using two-tailed unpaired t-test.

Next, we checked Ctt1 protein levels in Δ*scd6*. The Ctt1 protein levels are high in Δscd6 as compared to the WT in the untreated mid-log phase cells (Figure 1d, e) which decreases upon H_2_O_2_ treatment. There is no significant difference in the *CTT1* mRNA levels between WT and *Δscd6* (Figure S1d). The increased Ctt1 protein levels with the constant mRNA levels in Δ*scd6* suggests that there can be a post-transcriptional regulation.

### Δ*scd6* cells are more tolerant to H_2_O_2_ stress

The Ctt1 protein levels are increased upon the deletion of Scd6 in the unstressed condition (Figure 1e) but decrease upon 4mM H_2_O_2_ treatment. Therefore, we hypothesized that the increased Ctt1 protein levels under unstressed conditions might help the cells acclimatiseto the oxidative stress response. To test the physiological role of deletion of Scd6 in oxidative stress, we performed a colony forming unit (CFU) count assay for WT and Δ*scd6*. We observed 2.5 fold increased CFU count in *Δscd6*. The deletion of Scd6 made the cells more tolerant to oxidative stress induced by 4mM H_2_O_2_ (Figure 2a). On the contrary, the overexpression of Scd6 made the cells sensitive to oxidative stress (Figure 2b). This observation is consistent withthe decreased Ctt1 protein levels under the same condition (Figure 1a)

**Figure 2:**
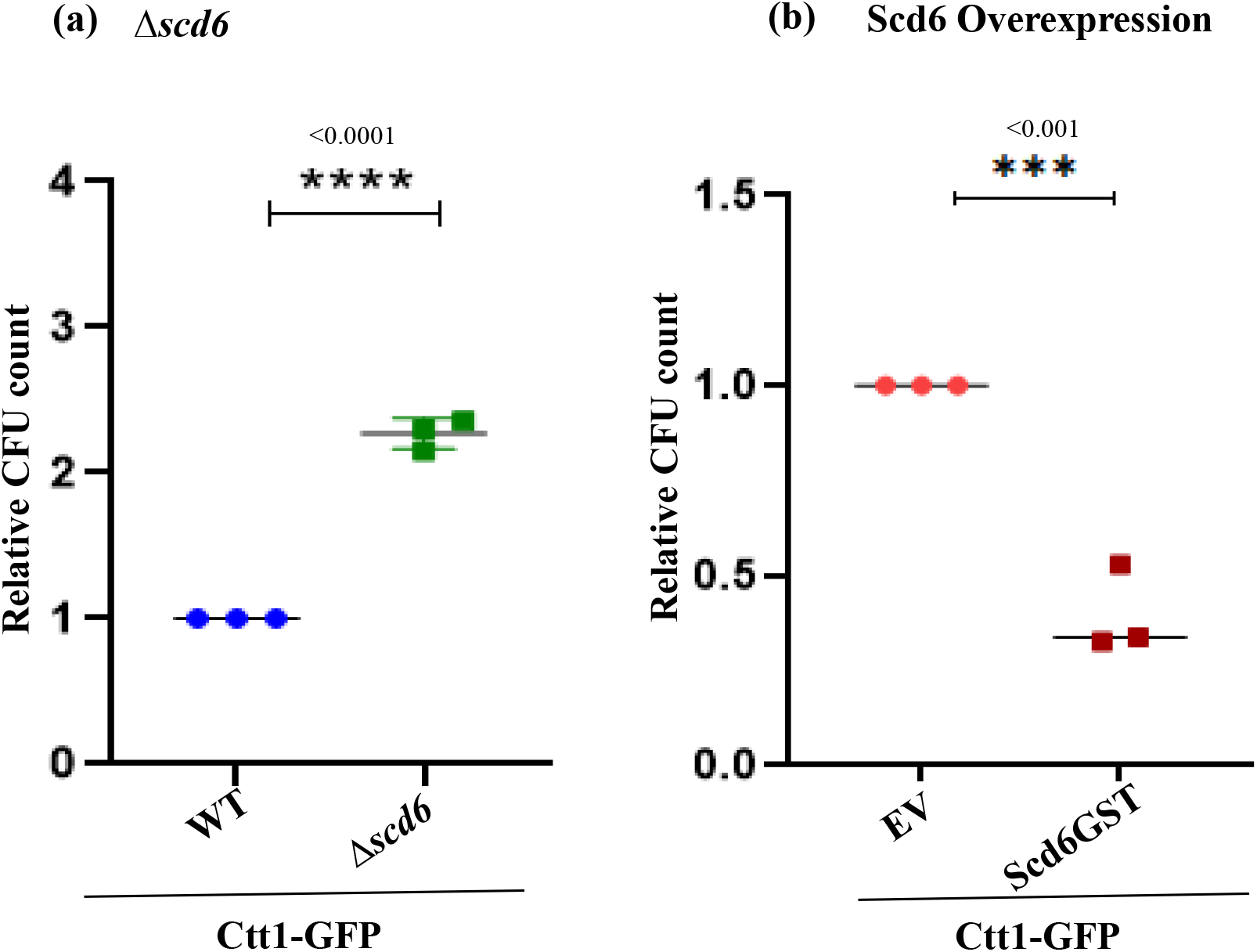
Alteration of Scd6 levels affects survival upon H_2_O_2_ stress. Mid-log phase Ctt1-GFP cells were treated with 4mM H_2_O_2_ for 30 minutes were plated and scored for their relative CFU count (Treated/untreated) a) CFU count assay showing the survival of Ctt1-GFP WT and *Δscd6* upon 4mM H_2_O_2_ treatment (b) CFU count assay showing the survival of Ctt1-GFP strain transformed with pYES EV and Scd6GST upon 4mM H_2_O_2_ treatment. ;*’ represents the statistical significance. It was calculated using two-tailed unpaired t test.

### Reactive oxygen species accumulate upon Scd6 overexpression in response to H_2_O_2_ treatment

The change in Ctt1 protein levels upon the change in Scd6 levels indicated that such changes could modulate ROS status of the cell. To test this, we performed DCFDA staining where we observed increased ROS accumulation in Scd6GST (H_2_O_2_+) as compared to EV and Scd6GST (Untreated) cells. The increased ROS accumulation upon Scd6 overexpression in response to H_2_O_2_ treatment can be one of the factors which canexplain the sensitivity of Ctt1-GFP Scd6GST on H_2_O_2_ treatment (Figure 2b). Although ROS levels in WT and *Δscd6* did not show a significant change in the ROS levels.

### Scd6 localizes to granules in response to H_2_O_2_ stress

Scd6 is known to localize to stress granules and P bodies upon glucose starvation and sodium azide stress^21,30^. Therefore, we tested the localization of Scd6 in response to H_2_O_2_ stress using live-cell imaging. Upon H_2_O_2_ treatment, these puncta decreases in number upon recovery indicating their dynamic nature (Figure 4a). To determine the identity of Scd6 puncta formed in response to H_2_O_2_ treatment, we checked the colocalization of Scd6 with Edc3 (core P body marker) and Pab1 (SG marker). Edc3 is known to form distinct puncta upon H_2_O_2_ treatment^5^. We observed an induction of Edc3mch puncta upon 4mM H_2_O_2_ treatment (Figure S3c). The co-localization experiments between Scd6 and Edc3 (a P-body marker) indicates that Scd6 localizes very poorly to P bodies upon H_2_O_2_ treatment (Figure S3d).To further test that Scd6 granules formed upon H_2_O_2_ treatment are stress granules, co-localization experiments were performed between Scd6 and the stress granule marker, Pab1. Live-cell imaging was performed in a Pab1RFP strain transformed with pRS316 Scd6GFP plasmid. The H_2_O_2_ treatment did not induce stress granules (Figure S3e) This is consistent with previous observations where H_2_O_2_ treatment did not induce stress granules^5^.

**Figure 3:**
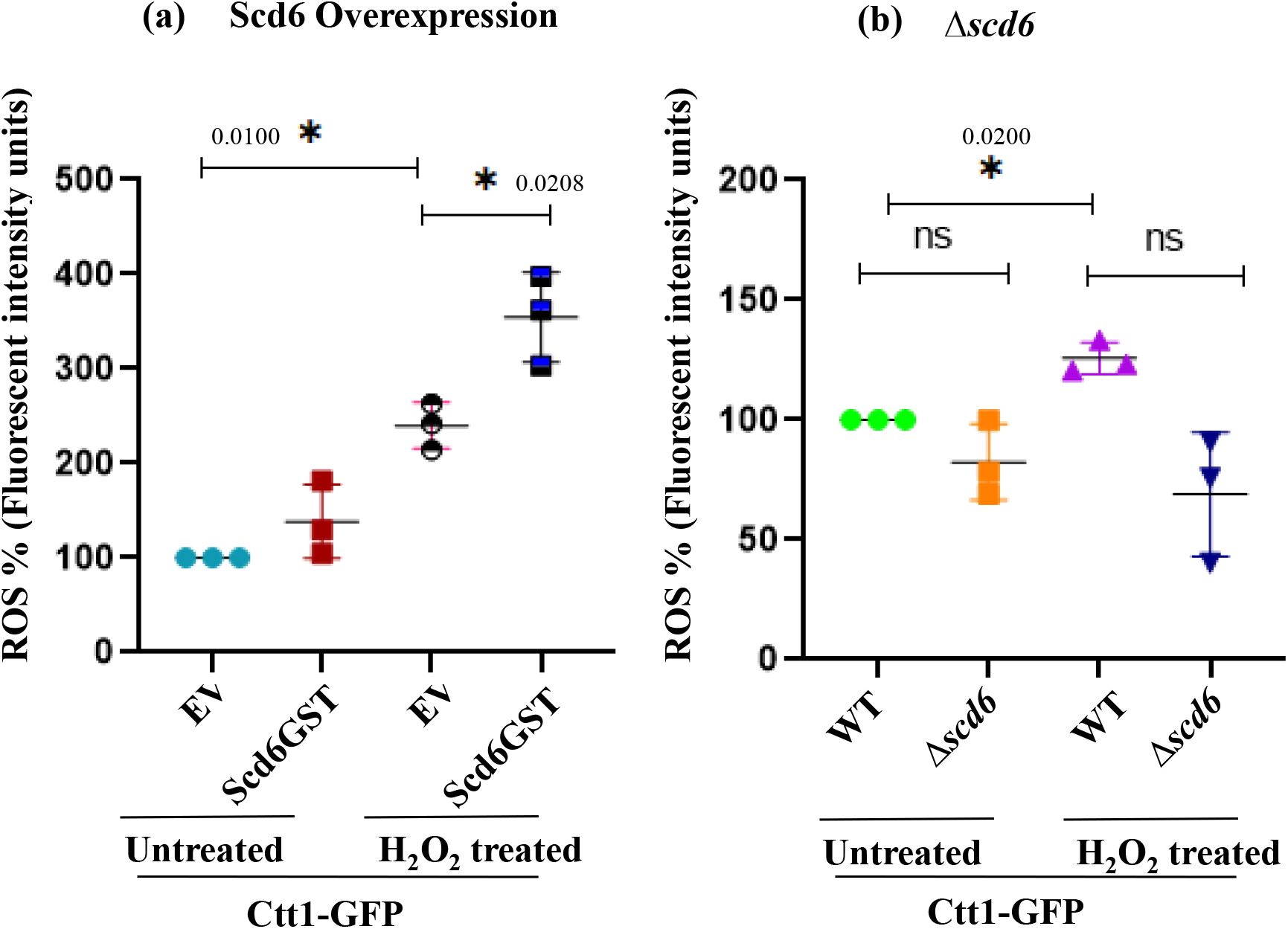
Alteration of Scd6 levels affects ROS levels: ROS detection using H_2_DCFDA. Quantitation for ROS % in (a) Scd6 overexpression and (b) *Δscd6* in untreated and H_2_O_2_ treated mid-log phase cells. ‘*’ represents the statistical significance. It calculated using unpaired Two-tailed student’s t-test.

**Figure 4:**
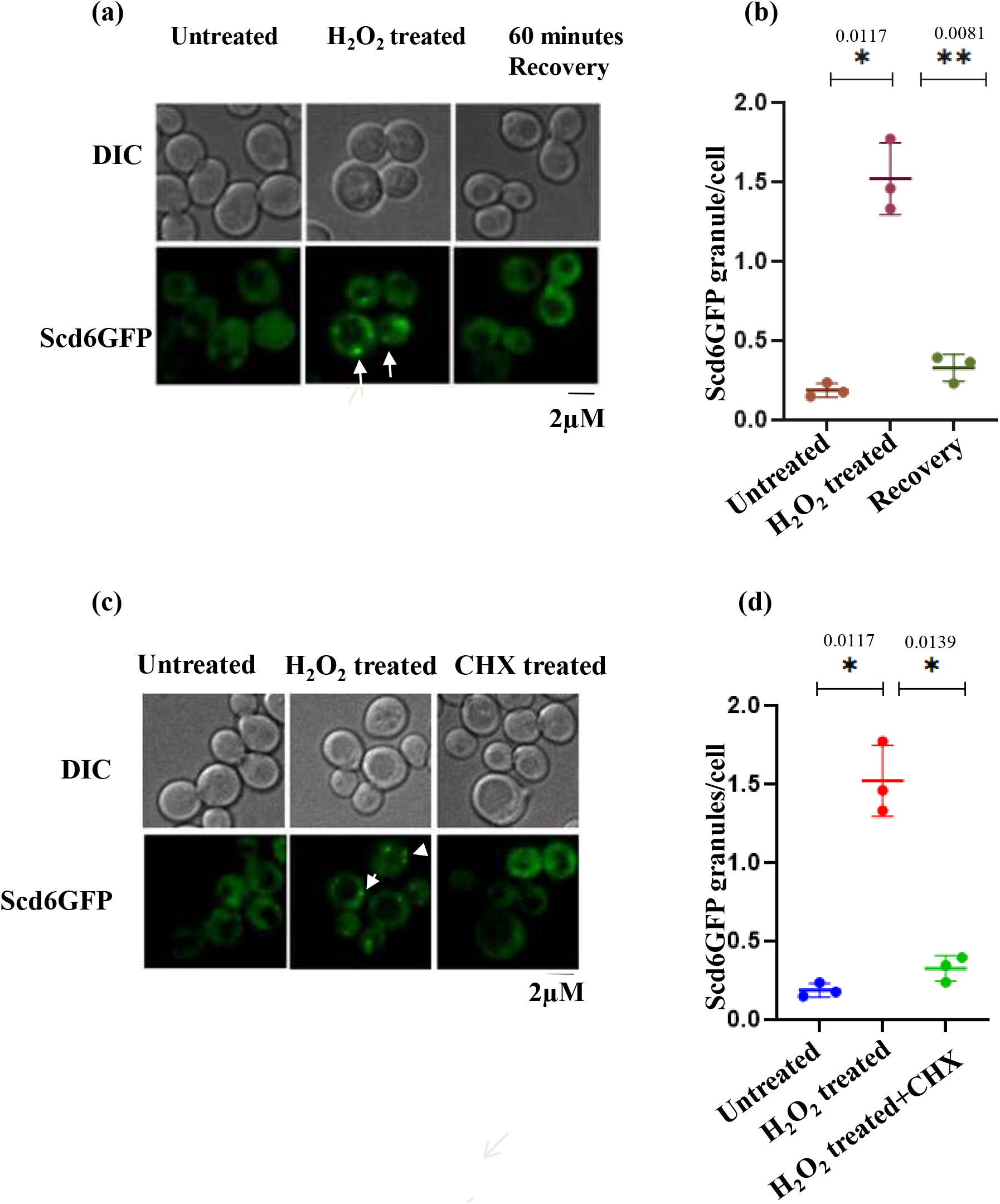
Scd6 forms puncta upon oxidative stress. (a) Live cell imaging of WT cells transformed with PRS316 Scd6GFP plasmid in untreated, 4mM H_2_O_2_ treated and Recovery conditions (b) Quantification for the Scd6GFP puncta formation in (a), (c) Live cell imaging for the localization of Scd6 upon 4mM H_2_O_2_ and CHX treatment (d) Quantification for Scd6GFP puncta formation showing upon 4mM H_2_O_2_ and CHX treatment ‘*’ represents the significance which was calculated using two-tailed paired t-test (n=3).

Scd6 localizes to puncta in response to H_2_O_2_which neither colocalizes with P bodies or stress granules (Figure S3).To understand whether these puncta contain mRNA, we checked the impactof cycloheximide treatmenton Scd6 granules.The Scd6 puncta formed in response to H_2_O_2_ treatmentdisappear in response to CHX treatment (Figure 4c,d)

### The methylation of Scd6 decreases upon H_2_O_2_ treatment

The Ctt1 protein levels are affected in a contrasting manner in untreated and treated condition (Figure 1a). In untreated cells, Scd6GST overexpression increases Ctt1 protein levels whereas upon H_2_O_2_ treatment, Ctt1 protein levels are decreased upon Scd6GST overexpression. PTMs like methylation have been shown to modulate the functions of RNA-binding proteins including Scd6. We hypothesised that methylation of Scd6 could be involved in modulating the behaviour of Scd6 in different conditions and therefore we tested the mono-methylation status of Scd6 upon H_2_O_2_stress treatment. We observed that the methylation level of Scd6 decreases upon H_2_O_2_ treatment (Figure 5).

**Figure 5:**
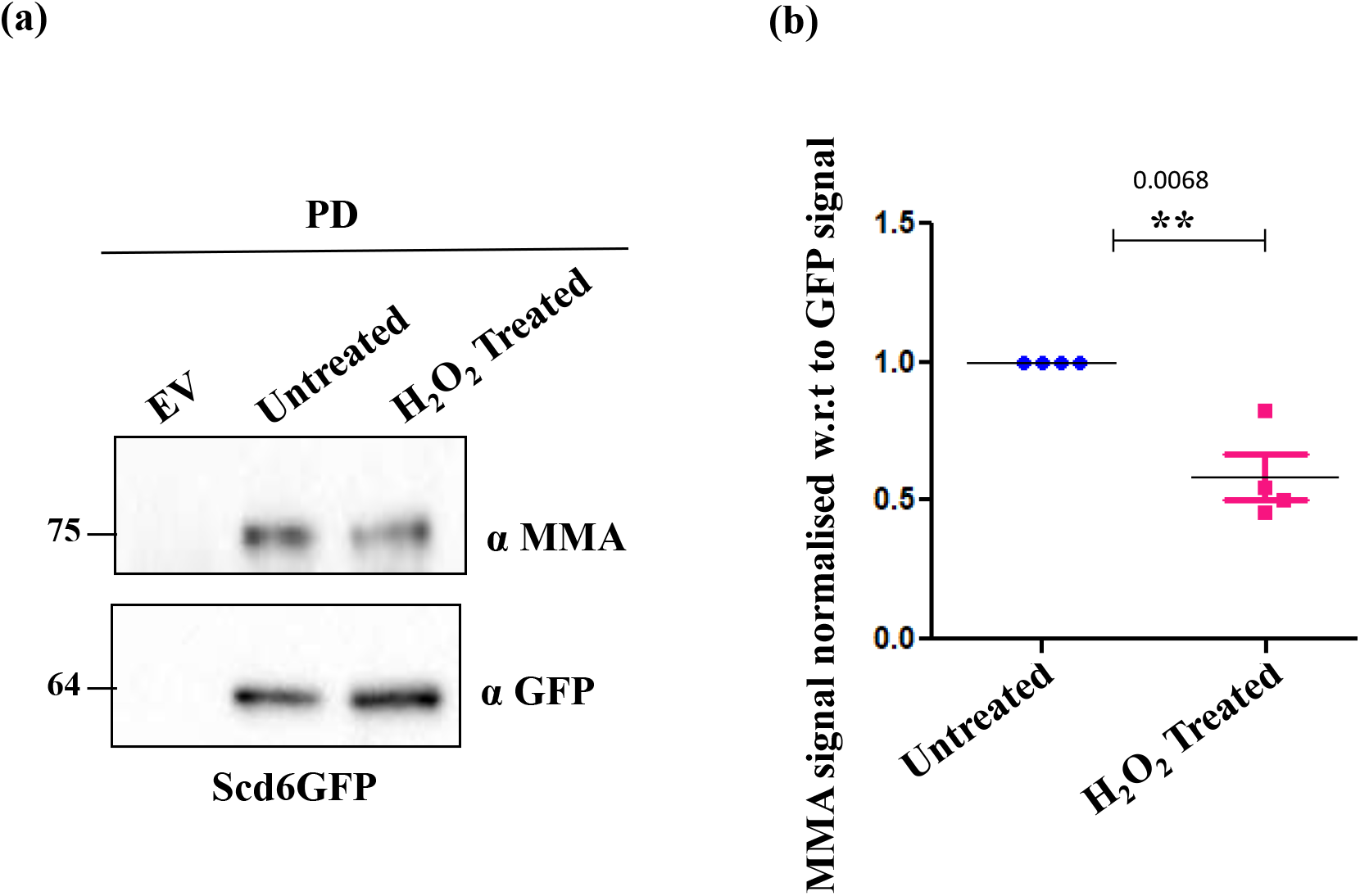
The methylation of Scd6 decreases upon H_2_O_2_ treatment. Scd6GFP was pulled down using a GFP-Trap followed by probing with mono Methyl Arginine (MMA) antibody and α-GFP (a) MMA levels of Scd6GFP in untreated and H_2_O_2_ treated condition (b) Quantification for the MMA signal normalised with GFP levels. ‘*’ represents the statistical significance. It calculated using two-tailed paired t-test.

### Scd6 interacts with *CTT1* mRNA

We tested whether Scd6 could bind *CTT1* mRNA to alter its translation. Scd6GST pulldown was performed for both untreated and H_2_O_2_ treated cells and RNA was isolated from the pull-down product followed by checking the enrichment of *CTT1* RNA using qRT PCR. *CTT1* was found to be significantly enriched in the overexpression of Scd6GST in untreated cells (Figure 6b).

**Figure 6:**
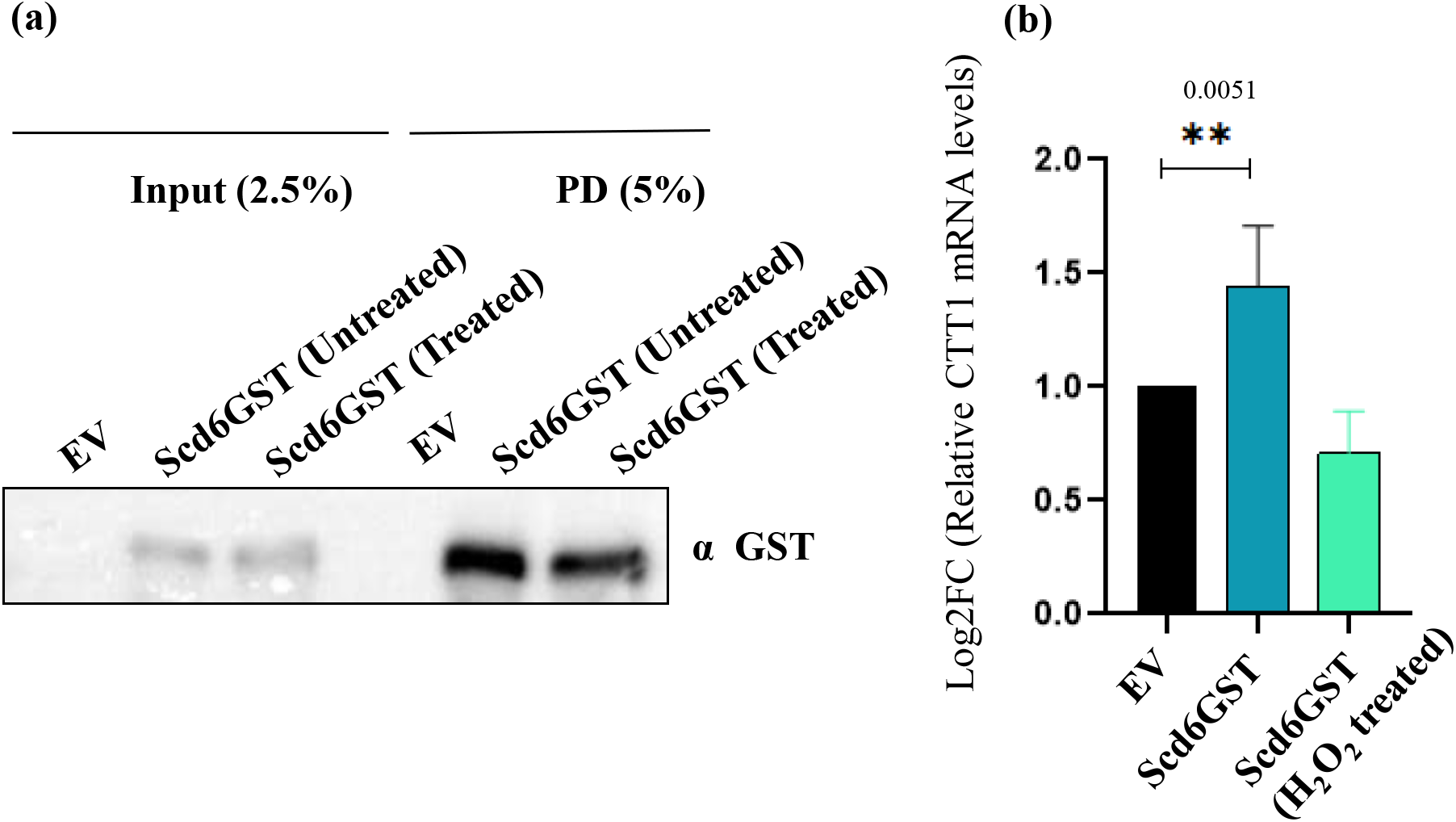
Scd6 binds *CTT1* mRNA (a) Blot represents Input and PD for Scd6GST pull-down. The GST signal was detected using α GST antibody (b) Log2FC for *CTT1* in the PD fraction showing the enrichment of *CTT1* mRNA in the Scd6GST pull-down in untreated and 4mM H_2_O_2_ treated condition. ‘*’ represents the statistical significance. It calculated using two-tailed unpaired t-test.

**Figure 7:**
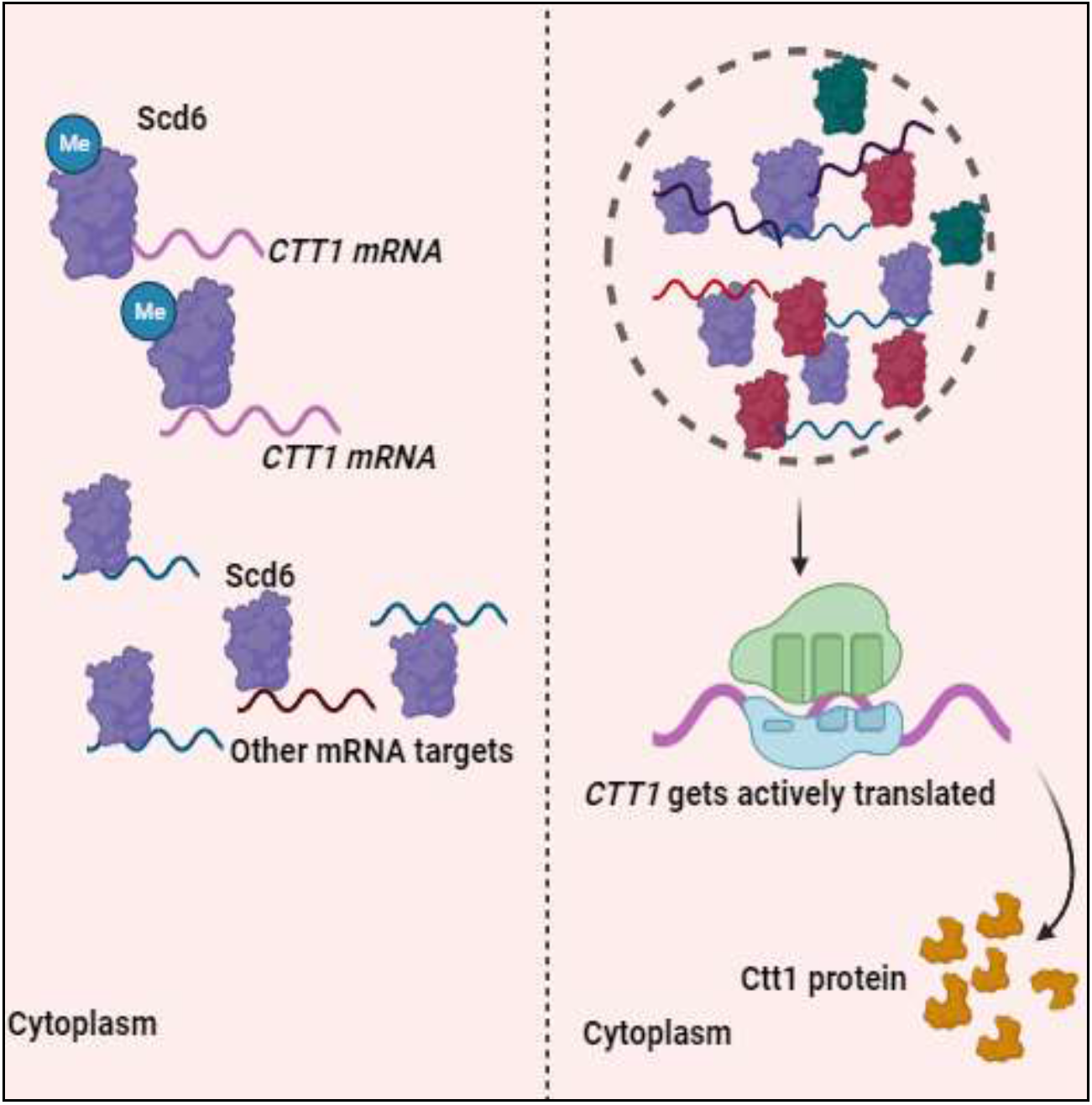
Model for the regulation of *CTT1* by Scd6: In the absence of any stress, the CTT1 mRNA serves as a target for the protein Scd6, forming a stable complex. However, when exposed to hydrogen peroxide (H_2_O_2_), there is a disruption of the *CTT1*-Scd6 interaction which could be due to a reduction in the methylation of Scd6, causing their dissociation. Under these oxidative stress conditions, Scd6 locates to a distinct puncta which is neither stress granule or p body within the cell. Meanwhile, the *CTT1* mRNA becomes actively engaged in translation in response to the H_2_O_2_ treatment. This interaction enables Scd6 to regulate the abundance of *CTT1* mRNA in untreated conditions, where it is not required in large quantities. In summary, H_2_O_2_ treatment induces the dissociation of *CTT1* and Scd6, leading to CTT1’s translation and Scd6’s localization to stress granules and P bodies.

## Discussion

In this study, we report the translational regulation of *CTT1* by of an RGG-motif containing protein, Scd6, the key observations supporting this finding are as follows: (1) Both in untreated and H_2_O_2_treated cells, we observed a significant alteration in Ctt1 protein levels when Scd6 was either overexpressed or deleted. Remarkably, these changes occurred without any corresponding alterations in mRNA levels (2) Scd6 overexpression rendered cells more susceptible to H_2_O_2_ treatment, whereas Scd6 deletion led to enhanced tolerance. (3) Cells accumulate more ROS upon Scd6 overexpression which makes them more sensitive (4) Scd6 localizes to puncta upon H_2_O_2_ treatment which are dynamic in nature (5) Scd6 binds *CTT1*mRNA.

Scd6 exhibits distinct effects on Ctt1 protein levels in a differential manner in untreated and H_2_O_2_ treated cells. In Scd6 overexpression, Ctt1 protein levels are increased in untreated cells whereas it decreases upon stress (Figure 1, a-c). The explanation for the differential behavior could be tied to the post-translational modifications of Scd6. Surprisingly, the deletion of Scd6 yields similar effects on Ctt1 protein levels (Figure 1,d-f) Even though the effect of Scd6 overexpression and deletion on Ctt1 protein levels are similar, the mechanism could be different. The increased Ctt1 protein level in *Δscd6* suggests an increase in translation of Ctt1 which is consistent with a previous observation^24^. However, in response to H_2_O_2_ treatment, Ctt1 protein levels begin to decrease. This phenomenon can be explained by considering that Δscd6 cells already possess a surplus of Ctt1 protein. When exposed to H_2_O_2_, the cellular demand for additional Ctt1 diminishes, leading to the reduction in Ctt1 protein levels in *Δscd6* cells.

The alteration in Ctt1 protein levels by Scd6 overexpression and deletion affects the survival of these cells in H_2_O_2_ induced oxidative stress. Scd6 overexpression makes the cells sensitive to H_2_O_2_ treatment whereas cells with scd6 deletion became tolerant to H_2_O_2_ treatment (Figure 2). This observation explains the direct impact of Scd6 on cellular responses to oxidative stress. The sensitivity and tolerance correlates well with the status of ROS in these cells. Cells with Scd6 overexpression which showed sensitivity to H_2_O_2_ treatment were found to have significantly increased ROS levels (Figure 3a). *Δscd6* cells with showed tolerance to H_2_O_2_ treatment did not show a significant change in ROS levels although there is a trend for decrease ROS (Figure 3b)

Scd6 localized to distinct puncta upon H_2_O_2_ treatment. These puncta decreases after the removal of stress which indicates they are dynamic in nature (Figure 4a). Colocalisation experiments suggest that these Scd6 puncta do not colocalise with Edc3 (P body marker) (Figure S3d). We did not observe any stress granule formation under similar conditions which is consistent with the previous osbervations^5^ (Figure S3e). Scd6 puncta formed upon H_2_O_2_ treatment are neither Pbody nor stress granules. Scd6 puncta formed upon H_2_O_2_ treatment decrease upon CHX treatment which indicates it contains RNA (Figure 4c, d)

Many RNA binding proteins are known to undergo post-translational modification. The affinity of RNA binding proteins towards their targets is factor which is often controlled by PTMs^31,32^. Scd6 is known to get methylated by Hmt1^22,33^. The methylation of Scd6 has been shown to be important for Scd6’s localization to P bodies^33^. In response to H_2_O_2_ treatment, the methylation of Scd6 decreases (Figure 5). We have also observed that Scd6 localizes to distinct puncta upon H_2_O_2_ treatment. Interestingly, *CTT1* interacts with Scd6 in untreated conditions (Figure 6b). This observation suggests a pre-existing interaction between Scd6 and *CTT1*, which may be integral to the regulation of *CTT1* expression. This pre-existing interaction may change depending upon the stress and need for Ctt1 in the cell. The decreased methylation in response to H_2_O_2_ treatment could act as a trigger to hamper the interaction between *CTT1* and Scd6 leading to the release of *CTT1* mRNA in the cytoplasm and concomitant localization of Scd6 to the distinct puncta which could be the regulatory sites of post-transcriptional control for another mRNA.

We propose that in untreated cells, the *CTT1* mRNA is present in an association with Scd6 (Figure 6b) which prevents its translation. However, when cells are exposed to H_2_O_2_ treatment, there is a dissociation of Scd6 from *CTT1* mRNA, leading to the localization of Scd6 to a distinct puncta (Figure 4a). The dissociation of Scd6 from *CTT1* mRNA could be a result of decreased methylation (Figure 5). Because of this dissociation, *CTT1* mRNA becomes actively engaged in translation in response to the H_2_O_2_ induced stress.Scd6 acts as a crucial regulatory factor that governs the translation of *CTT1* mRNA in response to oxidative stress.

## Supporting information

Supplementaryinformation_tiwarietal2023

## Notes

### Competing Interest Statement

The authors have declared no competing interest.

