## Supplementaryinformation_tiwarietal2023 for "RGG-motif protein Scd6 affects oxidative stress response by regulating Cytosolic caTalase T1 (Ctt1)"

### Supplementary figures

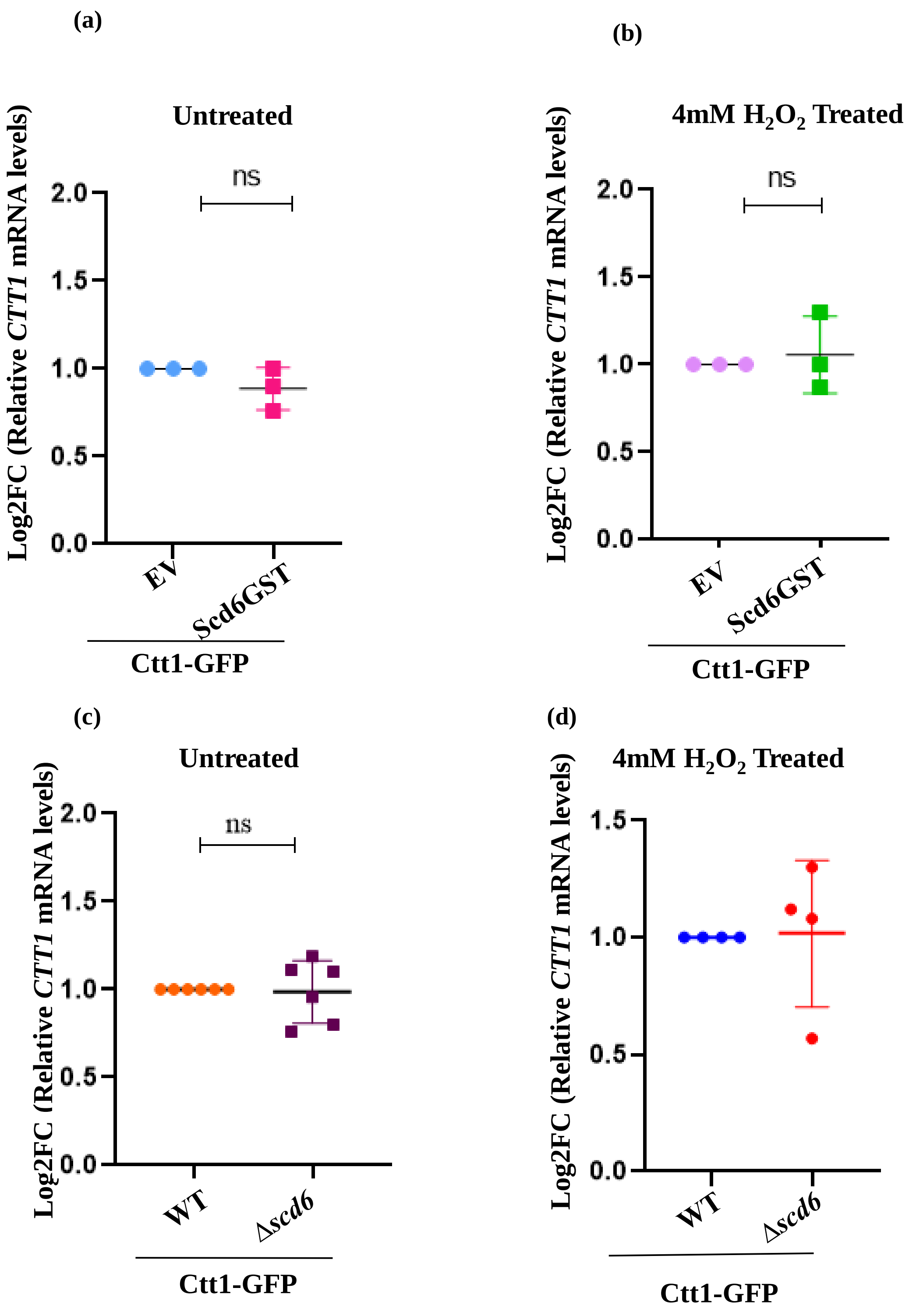


**Figure S1:** The *CTT1* mRNA levels remain constant upon Scd6 overexpression: (a) *CTT1* mRNA levels upon Scd6 overexpression (a) untreated and (b) 4mM $H_2O_2$ treated . *CTT1* mRNA levels upon Scd6 overexpression in Δ*scd6* (c) untreated (d) 4mM $H_2O_2$ treated. Significance was calculated using two-tailed unpaired –test.

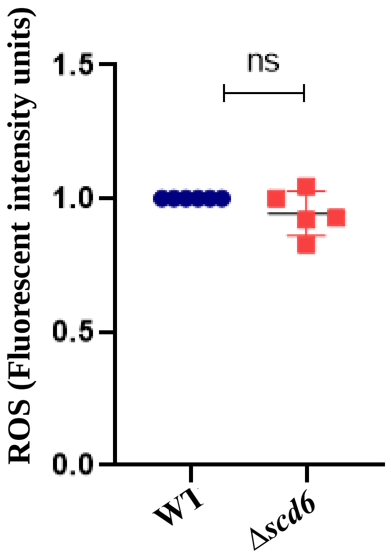


**Figure S2:** The ROS status does not change in Δ*scd6* (FACS)

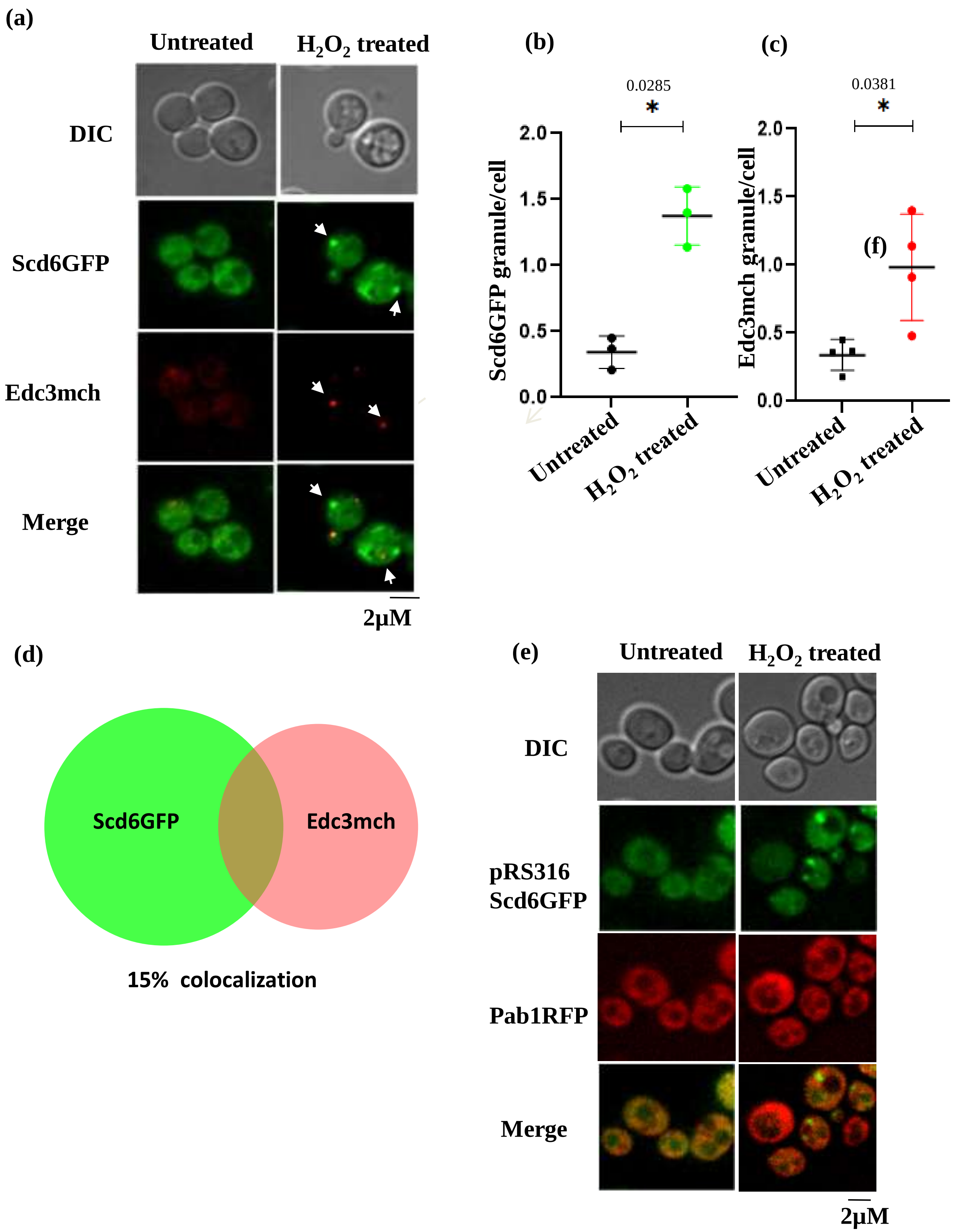


**Figure S3: : Scd6 puncta do not colocalize with Edc3 or Pab1**: (a) Live cell imaging of Scd6-GFP strain transformed with PRS416 Edc3mch in untreated, 4mM $H_2O_2$ treated condition (b) Quantification for Scd6GFP (c) Edc3mcherry puncta formation, (d) Venn diagram for the colocalization of Scd6GFP and Edc3mch (e) Live cell imaging of Pab1RFP cells transformed with PRS316 Scd6GFP plasmid in untreated, 4mM $H_2O_2$ treated condition.
